# A novel mechanism of bulk cytoplasmic transport by cortical dynein in *Drosophila* ovary

**DOI:** 10.1101/2021.11.12.468440

**Authors:** Wen Lu, Margot Lakonishok, Anna S. Serpinskaya, Vladimir I. Gelfand

## Abstract

Cytoplasmic dynein, a major minus-end directed microtubule motor, plays essential roles in eukaryotic cells. *Drosophila* oocyte growth is mainly dependent on the contribution of cytoplasmic contents from the interconnected sister cells, nurse cells. We have previously shown that cytoplasmic dynein is required for *Drosophila* oocyte growth, and assumed that it transports cargoes along microtubule tracks from nurse cells to the oocyte. Here we report that instead transporting cargoes along microtubules into the oocyte, cortical dynein actively moves microtubules in nurse cells and from nurse cells to the oocyte via the cytoplasmic bridges, the ring canals. We demonstrate this microtubule movement is sufficient to drag even inert cytoplasmic particles through the ring canals to the oocyte. Furthermore, replacing dynein with a minus-end directed plant kinesin linked to the actin cortex is sufficient for transporting organelles and cytoplasm to the oocyte and driving its growth. These experiments show that cortical dynein can perform bulk cytoplasmic transport by gliding microtubules along the cell cortex and through the ring canals to the oocyte. We propose that the dynein-driven microtubule flow could serve as a novel mode of cargo transport for fast cytoplasmic transfer to support rapid oocyte growth.

## Introduction

Microtubules perform many key cellular functions, such as cell division, migration, polarization/compartmentation, and intracellular long-range cargo transport. Cytoplasmic dynein (referred simply as dynein hereafter) is the major minus-end directed microtubule motor, and involved in numerous microtubule-based functions ^1^. In interphase cells, dynein is the main motor responsible for transporting various cargoes towards the microtubule minus-ends ^2^. In dividing cells, dynein functions at kinetochores, spindle poles, and at the cell cortex. Particularly, cortical dynein pulls astral microtubules and is therefore required for positioning the mitotic spindles ^3, 4^.

The core dynein complex contains two copies of heavy chain (DHC), intermediate chain (DIC), intermediate light chain (DLIC) and three different dynein light chains (Roadblock, LC8/Cut up and Tctex/Dlc90F) ^2, 5^ (Figure. 1A). Dynein heavy chain contains a ring of six AAA+ domains, and ATP hydrolysis-induced conformational change results in dynein walking towards the minus-ends of microtubules ^6^. The activity of dynein is regulated by the dynactin complex including the largest subunit p150^Glued^/DCTN1 ^7^, and the Lis1-NudE complex ^8^, as well as several activating adaptors, such as BICD2/BICDL1, Spindly and HOOK1/3 ^2, 5, 9^ (Figure. 1A).

**Figure 1.**
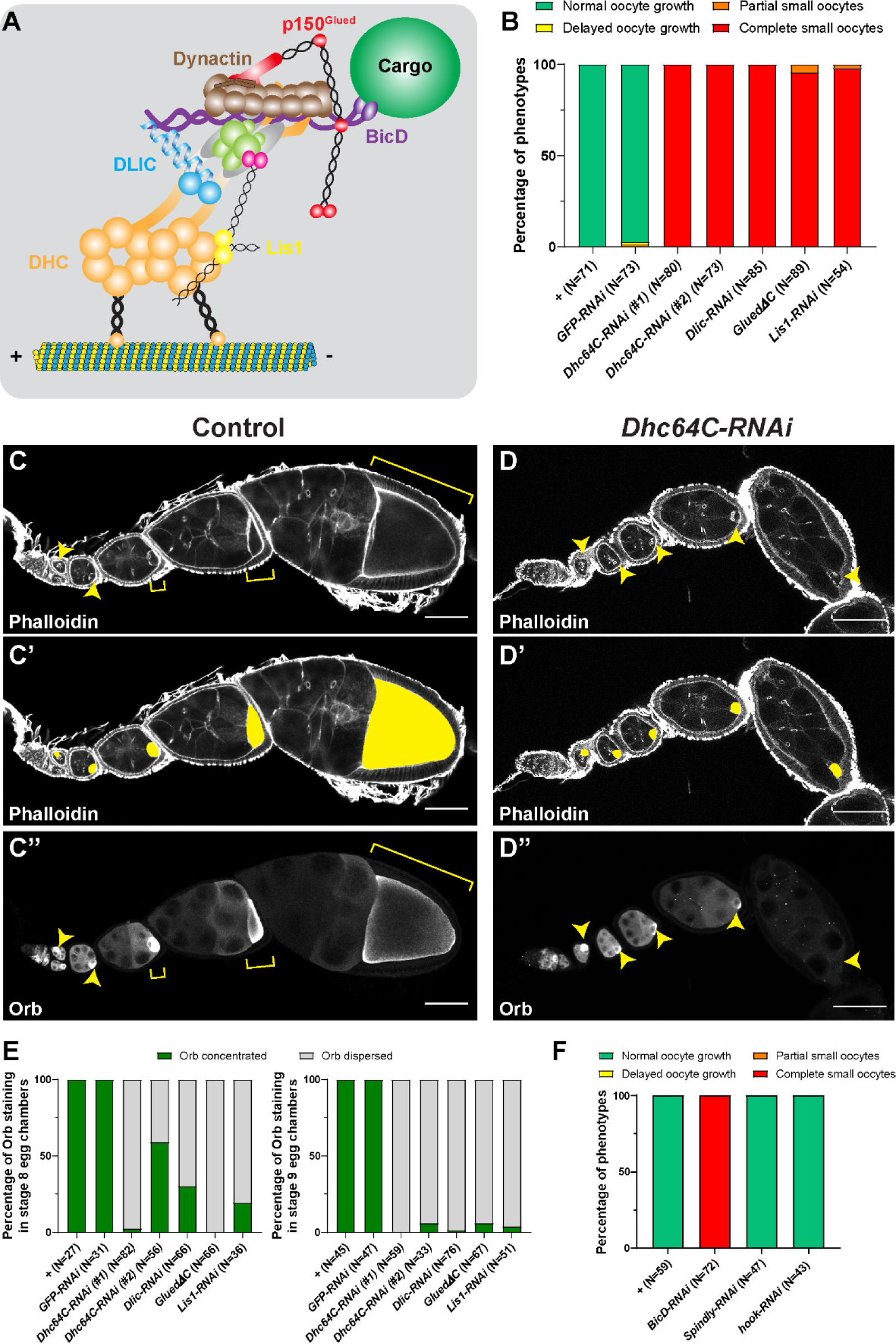
Dynein activity is required for *Drosophila* oocyte growth. (A) A cartoon illustration of cytoplasmic dynein and its regulators. The dynein core complex is composed of dimers of dynein heavy chain (orange), dynein intermediate chain (gray), dynein light intermediate chain (blue), and three types of dynein light chains (green). Dynein activity is regulated by the dynactin complex (brown, with p150^Glued^ highlighted in red) and the Lis1-NudE complex (Lis1, yellow; NudE, magenta). A dynein activating adaptor, BicD (purple), is also shown to illustrate the linkage of the dynein complex with a cargo. To note: other cargo adaptors instead of BicD, such as Spindly, HOOK1/3, ninein/ninein-like(NINL), and RAB11 family-interacting protein 3, can be used for dynein activation and cargo recruitment (not shown) ^2^. BICDR1 and HOOK3 could recruit two dyneins for increased force and speed (not shown) ^2^. (B) Summary of oocyte growth phenotypes in listed genetic background (all with one copy of *maternal αtub-Gal4^[V37]^*). Classifications of oocyte growth phenotypes are included in Supplementary Figure 1A. *Dhc64C-RNAi* (#1) is the RNAi line used for all *Dhc64C-RNAi* experiments in this study. (C-D’’) Phalloidin and Orb staining in control (C-C’’) and *Dhc64C-RNAi* (D-D’’) ovarioles. Oocytes and Orb staining are highlighted with either yellow arrowheads and brackets (C-D and C’’-D’’), or with yellow painting (C’-D’). Scale bars, 50 µm. (E) Summary of the Orb staining phenotypes in stage 8 (left) and stage 9 (right) egg chambers in listed genotypes (all with one copy of *maternal αtub-Gal4^[V37]^*). (F) Summary of oocyte growth phenotypes in *RNAi* lines against three listed dynein activating adaptors (all with one copy of *maternal αtub-Gal4^[V37]^*).

Dynein has a number of essential functions during *Drosophila* oogenesis. First, it is required for germline cell division and oocyte specification ^10, 11^. During mid-oogenesis, dynein is required for transport of mRNA ribonucleoproteins (RNPs) and organelles from nurse cells to the oocyte ^12–15^. Within the oocyte, dynein transports and anchors the anterior and dorsal determinants that are critical for axis determination for future embryos ^16–18^. During vitellogenesis, dynein in the oocyte regulates endocytic uptake and maturation of yolk proteins from the neighboring somatic follicle cells ^19^.

The *Drosophila* oocyte undergoes dramatic cell growth and polarization during oogenesis ^20^. Remarkably, the oocyte remains transcriptionally quiescent during most of the oogenesis. For its dramatic growth, the oocyte relies on its interconnected sister cells, nurse cells, for providing mRNAs, proteins, and organelles through intercellular cytoplasmic bridges called ring canals ^20, 21^. Previously, we showed that dynein heavy chain drives oocyte growth by supplying components to the growing oocyte ^15^. Here, we study the mechanism of dynein-dependent transport of cargoes from nurse cells to the oocyte. Surprisingly, we find that microtubules, which had previously been considered as static tracks for dynein ^12–15^, are robustly moved by dynein within the nurse cell cytoplasm, and more remarkably, from the nurse cell to the oocyte. We further demonstrate that this dynein-powered microtubule gliding creates cytoplasmic flow carrying cargoes in bulk to the oocyte, including neutral particles that do not interact with motors. Furthermore, we show that a gliding-only chimeric minus-end motor anchored to the cortex is sufficient to support transport of cargoes to the oocyte, and therefore the oocyte growth. Altogether, we describe here a novel mechanism for dynein-driven cytoplasmic transport: cortically-anchored dynein drives microtubule gliding, and microtubules in turn move cytoplasmic contents in nurse cells and through the ring canals, driving oocyte growth. This provides a fast and efficient mode of bulk transport of a wide variety of components supplied by nurse cells to the oocyte.

## Results

### Dynein is required for oocyte growth

The growing *Drosophila* oocyte is transcriptionally silent and mostly relies on its interconnected sister nurse cells for mRNAs, proteins, and organelles ^20, 21^. Dynein, as the main minus-end directed microtubule motor in *Drosophila*, has been implicated in nurse cell-to-oocyte transport ^12–15^.

Here we investigated the roles of the dynein complex and its regulators in oocyte growth by taking advantage of a germline-specific Gal4, maternal α tubulin-Gal4^[V37]^, that is expressed in germline cells after the completion of cell division and oocyte specification ^15, 22^. This approach bypasses the requirement for dynein in early oogenesis. Knockdown of dynein by expressing either RNAi against dynein components (DHC, DLIC, Lis1) or a dominant negative construct of p150^Glued^/DCTN1 (*p150^Glued^ΔC*)^12^ driven by this Gal4 line allowed for normal cell division and oocyte specification but caused complete arrest of oocyte growth (hereafter referred as the “small oocyte” phenotype) (Figure 1B-1D’’; Supplementary Figure 1A). In spite of the growth inhibition caused by dynein knockdown, we found that the oocyte marker, Orb (oo18 RNA-binding protein) ^23^ is properly concentrated in the early oocytes, emphasizing that our approach does not interfere with germline cell division or oocyte specification during early oogenesis. However, Orb is clearly dispersed from the small oocytes at later stages (stages 8-9) (Figure 1C’’-1D’’ and 1E;), implying defects of nurse cell-to-oocyte transport. These data demonstrate that dynein core components and regulators are indeed essential for oocyte growth, likely via transporting cargoes into the oocyte.

As dynein activity relies on various cargo activating adaptors, we knocked down the *Drosophila* homologs of three main dynein cargo-specific adaptors, BicD, Spindly, and Hook ^9, 24^ by RNAi in the germ line. Among the three adaptors, knockdown of BicD results in complete small oocytes, while *Spindly-RNAi* and *hook-RNAi* do not cause obvious oocyte growth defects (Figure 1F).

This lack of oocyte growth phenotype in *Spindly-RNAi* and *hook-RNAi* animals is not due to low efficiency of the RNAi lines themselves, as RNAi knockdown of Spindly and Hook display typical loss-of-function mutant phenotypes of these adaptors. Maternal knockdown of Spindly (*mat αtub[V37]>Spindly-RNAi*) caused all embryos to fail to hatch (N>200) and zygotic knockdown using a strong ubiquitous Gal4 (*Actin5C-Gal4*) led to 0% eclosion rate from *Spindly-RNAi* pupae (Supplementary Figure 1B) ^25^. Hook is not required for fly viability or fertility, but is required for proper bristle formation ^26^, and *hook-RNAi* animals phenocopied the classic hooked-bristle phenotype observed in *hook* null allele (*hook*^11^) (Supplementary Figure 1C-1E’) ^26^.

This set of data leads us to conclude that BicD is the most important dynein activating adaptor for *Drosophila* oocyte growth.

### Dynein drives microtubule gliding in nurse cells

Having established that dynein and its associated proteins are required for oocyte growth, we next examined dynein tracks, cytoplasmic microtubules. Microtubules are localized inside the intercellular cytoplasmic bridges, the ring canals (Figure 2A-A’’’’), consistent with previous reports ^12–15^. However, when we examined the dynamics of microtubules in live samples using photoconversion ^27–29^, we surprisingly discovered that microtubules in the nurse cells are not stationary; they move and snake around in the nurse cells (xFigure. 2B-2B’; Video 1). Robust microtubule movement in nurse cells can also be seen with fluorescently labeled microtubule-associated proteins (MAPs), EMTB-TagRFP ^30^, GFP-Patronin ^15, 30^, and Jupiter-GFP ^15, 27, 28, 30, 31^ (Videos 2-4). Even more remarkably, microtubules are seen moving from the nurse cell to the oocyte through the ring canal, which most robustly occurs in stage 9 egg chambers (Figure 2D-2F; Videos 2-4). Altogether, we conclude that microtubules are not static tracks for dynein; instead, they actively move within the nurse cells and from the nurse cell to the oocyte.

**Figure 2.**
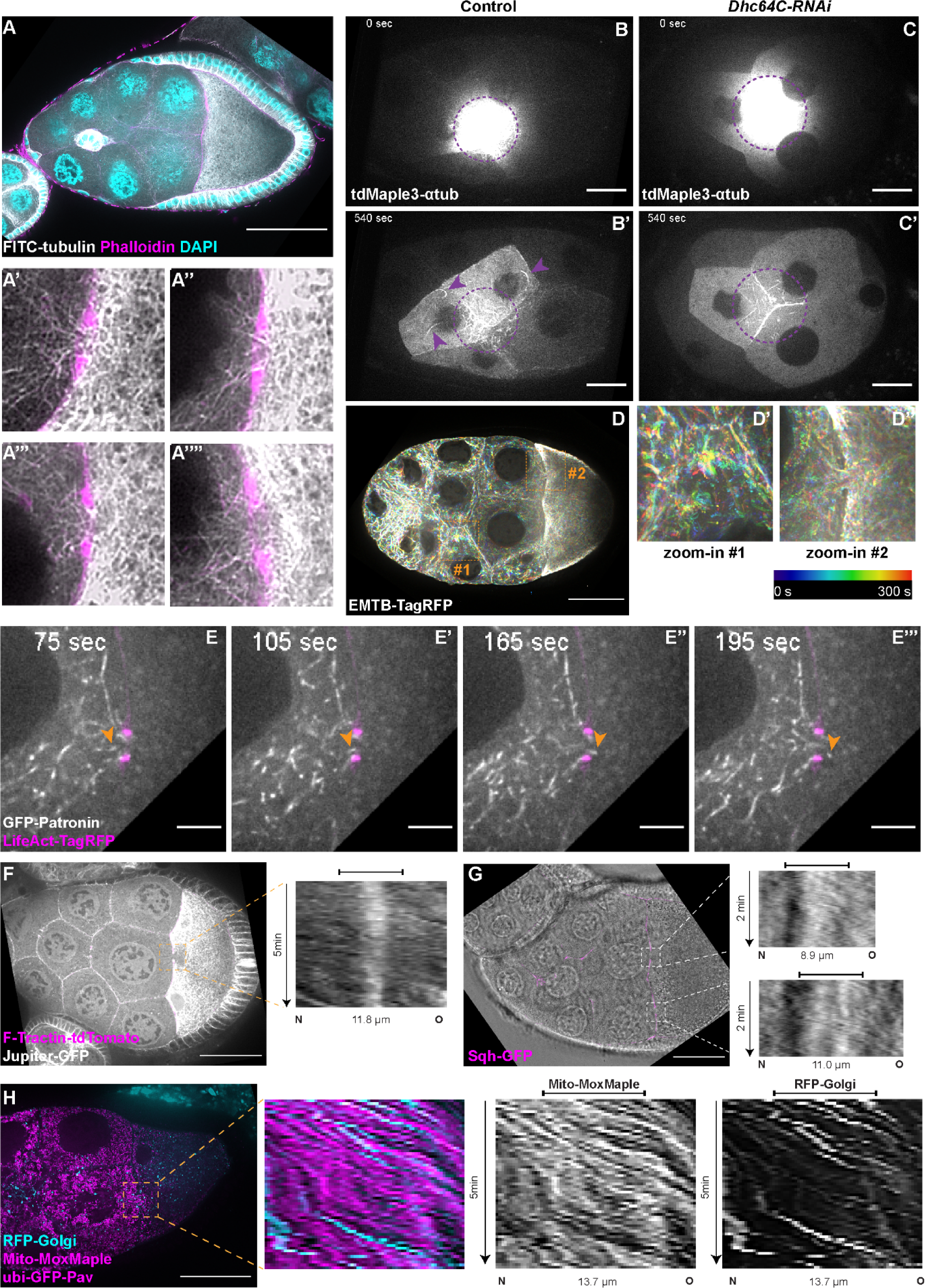
Microtubule movement within nurse cells and from nurse cells to the oocyte. (A) Tubulin staining in a control stage 9 egg chamber. Microtubules can be seen in all four ring canals connecting nurse cells with the oocyte (A’-A’’’’). Ring canals are labeled with rhodamine-conjugated phalloidin. Scale bar, 50 µm. (B-C’) Microtubule movement labeled with photoconverted tdMaple3-αtub in control and *Dhc64C-RNAi* nurse cells. Photoconversion area is highlighted with a dotted purple circle and microtubules outside of the photoconversion zone are highlighted with purple arrowheads. Scale bars, 20 µm. See also Videos 1 and 5. (D-D’’) Microtubule movement is visualized by a 3XTagRFP-tagged microtubule binding domain of human Ensconsin/MAP7 (EMTB-TagRFP). Temporal color-coded hyperstacks are used to show the microtubule movement in a whole egg chamber (D) and zoom-in areas of #1 within a nurse cell (D’) and #2 in a nurse cell-oocyte ring canal (D’’). Scale bar, 50 µm. See also Video 2. (E-E’’’) microtubule movement labeled with a GFP-tagged microtubule minus-end binding protein Patronin. The ring canal is labeled with LifeAct-TagRFP. One microtubule moving through the ring canal is highlighted with orange arrowheads. Scale bars, 10 µm. See also Video 3. (F) Microtubule movement is visualized by GFP protein trap line of an endogenous microtubule binding protein, Jupiter (Jupiter-GFP). The ring canal is labeled with F-Tractin-tdTomato. A kymograph of Jupiter-GFP in the nurse cell-oocyte ring canal (the dashed orange box) is used to show the microtubule movement from the nurse cell to the oocyte. The capped line on top of the kymograph indicates the ring canal region. Scale bar, 50 µm. See also Video 4. (G) Cytoplasmic flow is visualized by bright-field imaging. The ring canals are labeled with a GFP-tagged myosin-II light chain, Sqh-GFP. Kymographs of the two nurse cell-oocyte ring canals (the dashed white boxes) are used to show the cytoplasmic flow from the nurse cells to the oocyte. The capped lines on top of the kymographs indicate the ring canal regions. Scale bar, 50 µm. See also Video 7. (H) Bulk movement of two types of cargoes, mitochondria (magenta) and Golgi units (cyan), through the nurse cell-oocyte ring canal, labeled with GFP-Pav (magenta). A kymograph of mitochondria and Golgi units in the nurse cell-oocyte ring canal (the dashed orange box) is used to show that both cargoes move at a similar speed through the ring canal to the oocyte. The capped line on top of the kymograph indicates the ring canal region. Scale bar, 50 µm. See also Video 8.

When the *Drosophila* egg chambers reach stages 10B-11, nurse cells transfer all their cytoplasmic contents to the oocytes, the process called nurse cell dumping ^21, 32^. We examined whether the microtubule movement in the ring canals and associated oocyte growth during mid-oogenesis is a result of an early slow form of nurse cell dumping. We first measured the nurse cell size between stage 8 to stage 10 and found that nurse cells still undergo dramatic growth during this phase (Supplementary Figure 2A). This nurse cell growth from stage 8 to stage 10 is quite distinct from the dumping phase, during which nurse cells squeeze their cytoplasm to the oocyte and thus shrink quickly in size ^33^

Second, nurse cell dumping requires non-muscle myosin-II activity ^34, 35^. We tested whether myosin-II activity is required for stage 9 microtubule movement and overall oocyte growth. We used a RNAi line against the myosin-II heavy chain Zipper (*zip-RNAi*) to knockdown myosin-II activity, and an antibody recognizing phosphorylated myosin-II regulatory light chain (p-MRLC, Ser19) as a readout of myosin-II activity ^35, 36^. We found that that myosin-II activity is dramatically diminished in *zip-RNAi* egg chambers (Supplementary Figure 2B-2C’), suggesting that the *zip-RNAi* does inhibit myosin-II activity efficiently. However, we found that *zip-RNAi* does not inhibit microtubule movement in the nurse cell-oocyte ring canals (Video 6). Furthermore, *zip-RNAi* shows no defects in oocyte growth or Orb concentration from early to mid-oogenesis (Supplementary Figure 2D-2G). Additionally, we induced germline clones of a loss-of-function allele of *zip* (*zip*^2^)^37^, and found that the majority of *zip*^2^ mutant egg chambers with proper oocyte specification can develop to mid-oogenesis without obvious delay in oocyte growth (25 out of 28 egg chambers; Supplementary Figure 2H-2K). This is consistent with previous reports showing that the myosin-II light chain *spaghetti squash* (*sqh*) mutants ^34, 35^ and the “dumpless” mutants, such as chickadee, E2F, and RAP150B, develop normally to stage 10 without major oocyte growth defects ^38–40^. Thus, we conclude that the microtubule movement through ring canals is not a part of myosin-II-driven nurse cell dumping; it truly represents a novel process of nurse cell-to-oocyte transport in mid-oogenesis.

Dynein is known to glide microtubules *in vitro* and *in vivo* ^41^. Therefore, we asked whether dynein is the motor that moves microtubules in nurse cells. Knockdown of dynein using the *Dhc64C-RNAi* line results in a complete inhibition of microtubule movement within the nurse cells (Figure 2C-2C’; Video 5). The microtubules in *dynein-RNAi* nurse cells are noticeably less curved than the control ones, implying no motors applying forces on them. More importantly, the motility of microtubules in the nurse cell-oocyte ring canals are dramatically reduced in *dynein-RNAi* (Video 6). In contrast, we found that the microtubule movement through the ring canal is not affected by myosin-II inhibition (Video 6). Altogether these data support the idea that dynein glides microtubules in nurse cells and through ring canals to the oocyte.

### Microtubule flow carries cargoes to the oocyte

In addition to microtubule motility, we also observed cytoplasmic flow as well as synchronized movement of cargoes through nurse cell-to-oocyte ring canals in stage 9 egg chambers (Figure 2G-2H; Videos 7-8). Furthermore, cargo bulk movement is dynein-dependent and myosin-II-independent (Video 9). This raises the possibility that, in addition to the previously-proposed canonical cargo transport mode (Figure 3A), dynein-powered microtubule gliding could create cytoplasmic flow carrying all types of cargoes from nurse cells to the oocyte (Figure 3B).

**Figure 3.**
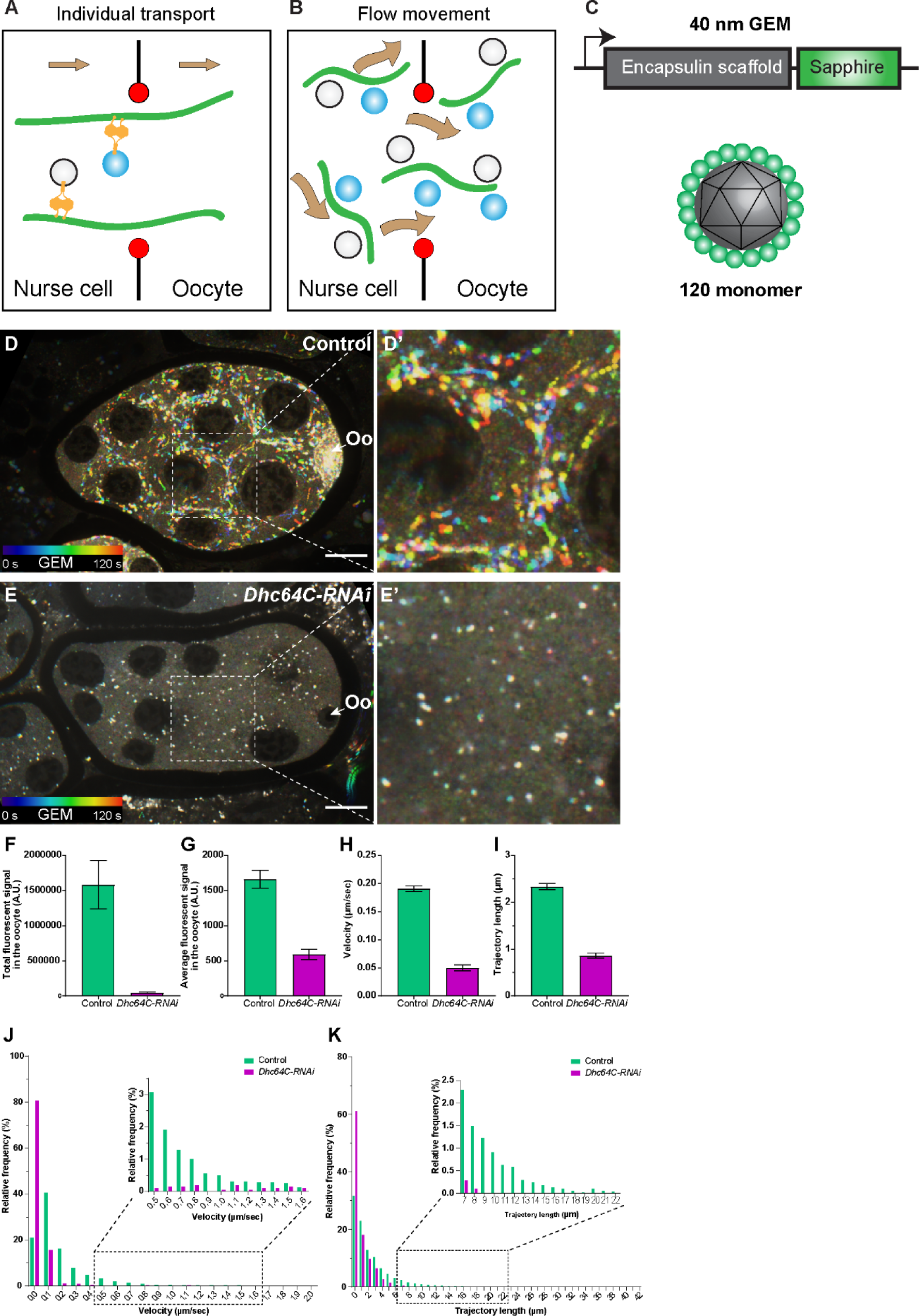
Dynein-dependent GEM particle movement in ovaries. (A-B) Cartoon illustrations of two possible mechanisms of dynein-dependent cargo transfer from the nurse cell to the oocyte. (C) A schematic illustration of the GEM construct. 120 copies of a sapphire-tagged *Pyrococcus furiosus* Encapsulin scaffold protein self-assemble into a 40 nm particle. (D-E’) Temporal color-coded hyperstacks of GEM particles in control (D-D’) and *Dhc64C-RNAi* (E-E’). Oo, the oocyte. Scale bars, 20 µm. See also Video 12. (F-G) Quantification of total (F) and average (G) fluorescent intensities of GEP particles in control and *Dhc64C-RNAi*. The values shown in the graphs are mean ± 95% confidence intervals. (F) Control, N=22; *Dhc64C-RNAi*, N=27. Unpaired t test with Welch’s correction between control and *Dhc64C-RNAi*: p<0.0001 (****). (G) Control, N=22; *Dhc64C-RNAi*, N=27. Unpaired t test with Welch’s correction between control and *Dhc64C-RNAi*: p<0.0001 (****). (H-K) Quantification of velocities (H, J) and trajectories (I, K) of GEM movement in control and *Dhc64C-RNAi*. The number of particles tracked: control, N=7656; *Dhc64C-RNAi*, N=2083. (H-I) The values shown in the graphs are mean ± 95% confidence intervals. (H) Unpaired t test with Welch’s correction between control and *Dhc64C-RNAi*: p<0.0001 (****). (I) Unpaired t test with Welch’s correction between control and *Dhc64C-RNAi*: p<0.0001 (****). (J-K) Histograms of velocities (J) and trajectories (K) of GEM movement in control and *Dhc64C-RNAi* (the same set of data used in H-I).

Therefore, to test this possibility, we simultaneously imaged both microtubules and organelles (mitochondria or Golgi units) in the nurse cell-oocyte ring canals, and found that they move together with microtubules through the ring canal to the oocyte (Videos 10-11). This suggests that mitochondria, Golgi units and probably other classes of cargoes could be carried through ring canals in bulk by microtubules gliding towards the oocyte (Figure 3B).

However, we cannot completely exclude the possibility that the motors attached to the cargoes walk on these moving microtubules and thus co-transport cargoes and microtubules through the ring canal. Thus, we decided to examine whether neutral cargoes that normally are not transported by motors along microtubules can also be transported to the oocyte. We used Genetically Encoded Multimeric nanoparticles (GEMs) (Figure 3C), which self-assemble into ∼40 nm fluorescent spheres ^42^. In *Drosophila* S2R+ cells, GEMs form bright compact particles and display mostly Brownian motion, which is very distinctive from typical movements of endogenous motor-driven organelles, such as lysosomes (Supplementary Figure 3). However, when we expressed GEMs in *Drosophila* ovaries, we found that they move within nurse cells in a fast linear manner (Figure 3D-D’; Video 12). More importantly, GEMs move through the nurse cell-oocyte ring canal (Video 13) and concentrate in the oocytes (Figure 3D and 3F-3G). Knockdown of dynein dramatically diminishes GEM linear movements and eliminates GEM accumulation in the oocyte (Figure 3E-3K; Video 12). Altogether, we demonstrate that the direct dynein-cargo interaction is not necessary for nurse cell-to-oocyte transport, and neutral particles can be efficiently carried through the ring canals by dynein-driven microtubule movement.

### Microtubule gliding delivers cargoes to oocytes

Next we investigated whether cortical dynein glides microtubules in nurse cells and transport cargoes to the oocyte. We first examined dynein localization using either a GFP-tagged Dlic transgenic line or an antibody against *Drosophila* dynein heavy chain and found a clear cortical localization of dynein in the nurse cells after the soluble pool of dynein is extracted by detergent (Supplementary Figure 4A-4D).

**Figure 4.**
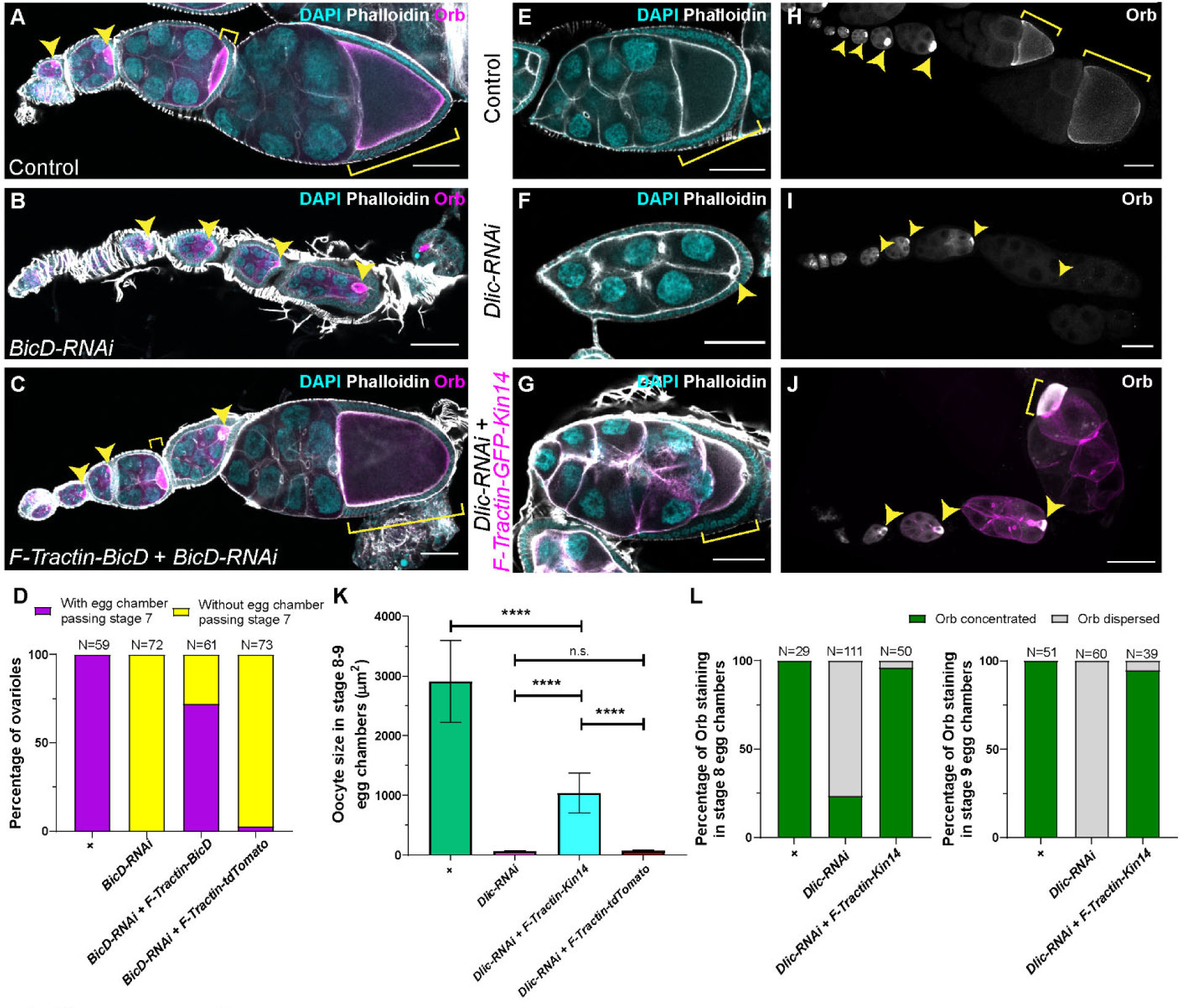
A cortically-anchored minus-end motor is sufficient for oocyte growth. (A-C) Oocyte growth defect in *BicD-RNAi* is rescued by a cortically-restricted BicD construct. (D) Summary of percentages of ovarioles with and without egg chamber(s) passing stage 7. (E-J) The defects of oocyte growth and Orb concentration in *Dlic-RNAi* are rescued by a cortically recruited plant kin14 (Kin14VIb) construct. Phalloidin and DAPI staining (E-G) or Orb staining (H-J) in control (E, H), *Dlic-RNAi* (F, I) and *Dlic-RNAi* rescued by F-tractin-GFP-Kin14 (G, J). (K) Quantification of oocyte size in stage 8∼9 egg chambers in listed genotypes. The values shown in the graph are mean ± 95% confidence intervals. Control, N=40; *Dlic-RNAi*, N=49; *Dlic-RNAi + F-Tractin-Kin14*, N=33; *Dlic-RNAi + F-Tractin-tdTomato*, N=42. Unpaired t test with Welch’s correction were performed in following groups: between control and *Dlic-RNAi*, p<0.0001 (****); between Control and *Dlic-RNAi + F-Tractin-Kin14*, p<0.0001 (****); between Control and *Dlic-RNAi + F-Tractin-tdTomato,* p<0.0001 (****); between *Dlic-RNAi* and *Dlic-RNAi + F-Tractin-Kin14*, p<0.0001 (****); between *Dlic-RNAi* and *Dlic-RNAi + F-Tractin-tdTomato,* p=0.2107 (n.s.); between *Dlic-RNAi + F-Tractin-Kin14* and *Dlic-RNAi + F-Tractin-tdTomato,* p<0.0001 (****). (L) Summary of Orb staining phenotypes in stage 8 (left) and stage 9 (right) egg chambers in listed genotypes. Oocytes are highlighted with either yellow arrowheads or yellow brackets. All listed genotypes carried one copy of *maternal αtub-Gal4^[V37]^*. Scale bars, 50µm.

Then we tested whether a cortically-anchored minus-end-directed motor is sufficient to drive oocyte growth. To constrain dynein activity to cell cortex, we replaced the endogenous dynein activating adaptor BicD with an ectopically expressed BicD that is cortically-recruited by an actin-targeting motif, F-Tractin ^43^ (Supplementary Figure 4E-F’). Compared to *BicD-RNAi* in which no egg chambers developed passing stage 7, this cortically-recruited BicD construct allows >70% ovarioles to have egg chambers reaching mid-oogenesis with proper Orb concentration (Figure 4A-4D). In contrast, expression of a tdTomato-tagged F-Tractin alone ^43^ (not fused with BicD) did not rescue the oocyte growth defects in *BicD-RNAi* (Figure 4D), indicating that the rescue we observed with F-Tractin-BicD is not due to Gal4 activity dilution or F-Tractin overexpression.

To further test the idea that gliding microtubules drive nurse cell-to-oocyte transport, we created an artificial minus-end gliding-only motor. It contains a dimer motor region of a fast minus-end-directed plant kinesin-14, kin14Vib ^44–46^, and is targeted to cell cortex with the F-Tractin probe (Supplementary Figure 4E). This chimeric kinesin-14 motor, unlike dynein, cannot carry endogenous *Drosophila* cargoes, allowing us to test whether microtubule gliding alone is sufficient for moving organelles to the oocyte and support the oocyte growth. With this chimeric gliding-only kinesin-14 motor, we found that oocyte growth is partially rescued (Figure 4E-4G and 4K). In addition to the oocyte size rescue, we also found that in more than 95% of the samples, the kinesin-14 motor is able to maintain Orb concentration in the oocyte of stage 8∼9 egg chambers (Figure 4H-4J and 4L). The rescues of both oocyte growth and oocyte Orb concentration imply that the gliding-only motor restores the nurse cell-to-oocyte transport.

Therefore, we directly examined mitochondria movement in the kinesin-14 rescued samples. We observed highly motile mitochondria in nurse cells, and synchronized mitochondria movement from the nurse cells to the oocyte, highly resembling mitochondrial flow in control ring canals (Video 14).

In summary, we show that cortically-anchored microtubule minus-end motors that cannot directly transport cargoes drive oocyte growth, further supporting our model that dynein-driven microtubule gliding powers cytoplasmic flow from the nurse cells to the oocyte (Figure 5; Video 15).

**Figure 5.**
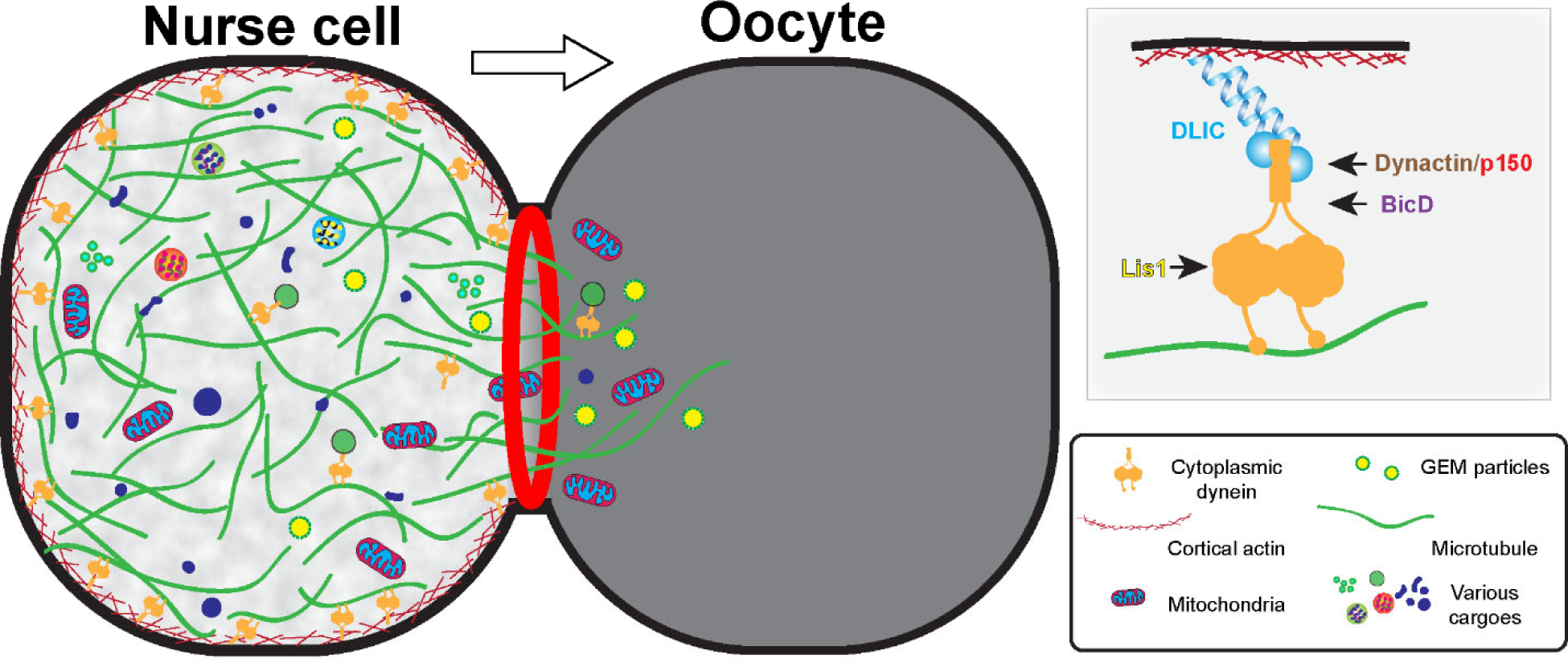
The model of cortically anchored dynein transferring cytoplasmic contents to the growing oocyte via gliding microtubules. Dynein light intermediate chain (Dlic) bridges dynein heavy chain to the cell cortex, while the Dynactin/p150 complex, BicD and Lis1 are required for activation of cytoplasmic dynein. In addition to the bulk cytoplasmic flow, dynein-dependent individual cargo transport on microtubules may also contribute to the nurse cell-to-oocyte transfer and oocyte growth. See also Video 15.

### C-terminus of dynein light intermediate chain is sufficient for nurse cell cortical localization

Having established that cortically anchored dynein is the key to glide microtubules and deliver cargoes to the growing oocyte, we decided to investigate how dynein is anchored to the nurse cell cortex. As we observed the cortical localization of Dlic-GFP in nurse cells (Supplementary Figure 4B), we decided to make N-terminal and C-terminal truncations of Dlic (DlicNT and DlicCT; Supplementary Figure 5A) and examine their localizations in the germ line. The Dlic N-terminus carries a GTPase-like domain and is known to interact with dynein heavy chain via a patch of conserved aromatic residues ^47^. The C-terminal Dlic contains the effector-binding domain that interact with BICD, Spindly and Hook-family activating adaptors to form a stable processive dynein-dynactin complex ^24^. We tagged the DlicNT and DlicCT with an optogenetic system LOVTRAP ^48^, and in dark LOVTRAP probes bring the two Dlic truncations together (Supplementary Figure 5A-5B). We found that the DlicNT-DlicCT complex in dark is sufficient to rescue the oocyte growth defects caused by *Dlic-RNAi* (Supplementary Figure 5C), indicating that the Dlic truncations are functional in the germ line. While DlicNT appears diffused in germline cytoplasm, DlicCT shows a strong cortical localization in the nurse cells (Supplementary Figure 5D-5E).

Since the DlicCT is known to interact with dynein activating adaptors [47] and BicD is the most important activating adaptor for oocyte growth (Figure 1F), we then tested whether BicD is required for the localization of DlicCT to the nurse cell cortex. In *BicD-RNAi* background, DlicCT still localizes to the cortex and mostly evidently at the ring canal regions (Supplementary Figure 5F), indicating that BicD is not essential for recruiting Dlic to the nurse cell cortex.

The dynactin complex includes a short actin-like filament composed of actin related proteins (Arp1 and Arp11) and β-actin that can potentially links dynein to the cell cortex. Previous studies in mitosis revealed that the dynactin complex facilitates dynein localization at the cell cortex for spindle pulling and positioning ^49, 50^. Therefore, we examined the DlicCT localization in the dominant negative p150^Glued^/DCTN1 mutant (*p150^Glued^ΔC*) and found that the DlicCT localization is still predominantly cortical with the inhibition of the dynactin complex (Supplementary Figure 5G). On the other hand, Lis1 has been shown to localize at the oocyte cortex and it is required to recruit dynein to the oocyte cortex ^51^. However, we found that the cortical localization of DlicCT in nurse cells is not affected by *Lis1-RNAi* (Supplementary Figure 5H).

In conclusion, Dlic C-terminus could facilitate the dynein complex to localize to the nurse cell cortex, independent of the dynein activating components (BicD, dynactin and Lis1). We propose that Dlic links the dynein heavy chain to the cell cortex and thus essential for microtubule gliding in nurse cells (Figure 5).

## Discussion

As the main microtubule minus-end directed motor in animal cells, cytoplasmic dynein is responsible for a wide variety of cellular functions, ranging from cell division to intracellular transport. In *Drosophila* ovary, dynein plays an essential role in nurse cell-to-oocyte transport of mRNAs and organelles ^12–15^. This was logically attributed to the conventional mode of dynein-driven transport: the motor attached to the cargo moves on microtubule tracks located inside ring canals and carries cargoes to the oocyte (Figure 3A).

In this study, we reveal a novel mechanism of bulk cargo transport by cytoplasmic dynein. First, we show that dynein core components and its regulatory cofactors are required for *Drosophila* oocyte growth (Figure 1). By imaging microtubules in live ovaries, we demonstrate that microtubules are actively moved by dynein from nurse cells to the growing oocytes (Figure 2). Furthermore, we use an artificial cargo that does not bind motors and show that direct dynein-cargo interaction is not necessary for the cargo movement in nurse cells or its transporting to the oocyte (Figure 3), supporting a passive “go-with-the-flow” mechanism underlying cytoplasm transfer from nurse cells to the oocyte. Lastly, we build a chimeric gliding-only motor by anchoring a minus-end plant kinesin, kin14VIb to the cortex, and find that this chimeric motor is sufficient to drive organelle transport and oocyte growth (Figure 4). Therefore, we propose a novel mechanism of dynein for bulk cargo transport: cortically-anchored dynein glides microtubules in the nurse cells; in turn these gliding microtubules move cytoplasmic contents within the nurse cells and from the nurse cells to the oocyte through the ring canals (Figure 5; Video 15).

### A novel phase of nurse cell-to-oocyte transport

Previously, nurse cell-to-oocyte transport has been divided into two phases: the early slow selective phase and the late fast non-selective phase ^21, 32^. The early phase is characterized by dynein-driven cargo transport along microtubules to the oocyte ^12–15^. The late massive nurse cell-to-oocyte transport phase is known as nurse cell dumping, which occurs at late stage 10B to stage 11 ^21, 32^.

Here we report a new phase of cytoplasmic flow driven by microtubules that are transported by cortical dynein from nurse cells to the oocyte. This occurs between the two previously-described phases, slow selective transport and fast nurse dumping. As microtubules can drag adjacent contents in the viscous cytoplasm ^29, 52^, we believe that this transport is non-selective. This conclusion is supported by the fact that the neutral particles (GEMs) are moved in the nurse cells and concentrated in the oocyte by dynein. Furthermore, we have found that a chimeric gliding-only motor unrelated to dynein, kinesin-14, is able to rescue mitochondria transport from the nurse cell to the oocyte (Video 14). As we use a motor domain of a moss kinesin-14 that has no known homology with the motor proteins that interact the mitochondrial adaptor protein Milton (e.g., KHC and Myosin 10A) ^53, 54^, we conclude that the flow created by microtubule movement in the ring canals is not cargo-specific, and it is different from the early selective transport phase.

During the dumping phase, nurse cells “squeeze” all the cytoplasmic contents to the oocyte, which is caused by non-muscle myosin-II contraction ^33, 35^, and associated with fast cytoplasmic streaming occurring in the oocyte mixing the dumped contents with the ooplasm ^21, 52, 55^. We reasone that the dynein-driven microtubule flow we observed is distinct from nurse cell dumping: (1) it occurs prior to nurse cell dumping and ooplasmic streaming (stages 8∼9 versus stages 10B-11); (2) the nurse cell size grows drastically during microtubule flow stages (Supplementary Figure 2A), instead of fast shrinking associated with nurse cell dumping; (3) it requires cytoplasmic dynein, instead of myosin-II activity (Supplementary Figure 2; Videos 6 and 9); (4) the “dumpless” mutants develop normally without major oocyte growth defects to stage 10 ^38–40^, which is noticeably different from the small oocyte phenotype we observed in dynein knockdown (Figure 1).

Therefore, the dynein-microtubule driven bulk cargo flow from nurse cells to the oocyte presents a novel phase between early selective transport and late non-selective dumping, which is essential for *Drosophila* oocyte growth.

### Dynein anchorage at the cell cortex

In this study, we found that cortical dynein glides microtubules to create local cytoplasmic flow and move cargoes to the growing oocyte. Furthermore, we found that the C-terminus of dynein light intermediate chain (DlicCT) is sufficient to target to the nurse cell cortex. DlicCT contains the effector-binding domain that interacts with multiple dynein activating adaptors ^24^. Recently, our lab reported that Spindly, a dynein activating adaptor that interacts with DlicCT ^24^ and recruits dynein to the kinetochore in mitosis ^25^, anchors dynein to cortical actin in axons, thus pushing microtubules of the wrong polarity out of the axons in *Drosophila* neurons ^56^. However, knockdown of Spindly does not disrupt dynein-dependent oocyte growth (Figure 1F). Thus, it indicates that the ovary uses a different mechanism for dynein cortical targeting. We showed that cortically-recruited BicD is able to rescue the oocyte growth arrest caused by *BicD-RNAi* (Figure 4A-4D), implying that BicD could contribute to the linkage between dynein and the cortex. Other studies also have suggested that the dynactin complex and Lis1 are involved in dynein cortical anchorage ^49–51^. Nevertheless, DlicCT cortical localization is unaffected after inhibition of BicD, dynactin/p150 or Lis1 (Supplementary Figure 5E-5H), again indicating a novel mechanism of anchoring dynein to the cortex in nurse cells. Intriguingly, Dlic physically interacts with the *Drosophila* Par-3 homolog, Bazooka (Baz), in the ovary ^57^, and Baz is a known cortical landmark protein in ovary and bind other cortical proteins, such as β-catenin (Armadillo) and DE-Cadherin (Shotgun) ^58–60^. Thus, it is tempting to propose that Bazooka is the key to anchor the dynein complex to the nurse cell cortex via its interaction with Dlic and thus allows dynein to glide microtubules along the cortex.

We consistently observe cytoplasmic flow from the nurse cells to the oocyte in stage 9 egg chambers, suggesting a direction-controlling mechanism underlying the persistent oocyte growth. We speculate that the flow directionality could be attributed to different levels of dynein gliding activity. The dynein complex is strongly localized to the nurse cell cortex, but to a much lesser extent to the oocyte cortex (Supplementary Figure 4). Consistent with the dynein localization, microtubules are more cortically localized in the nurse cells than in the oocyte (Figure 2F). This differences in dynein localization and microtubule organization may result in a higher dynein-driven microtubule gliding activity in the nurse cells, and therefore creates the directional flow through the ring canals to the growing oocyte.

Alternatively, this directionality could be controlled by the gatekeeper protein Short stop (Shot). Recently, we demonstrated that Shot is asymmetrically localized at actin fibers of ring canals on the nurse cell side and controls the cargo transport directionality between nurse cells and the oocyte ^15^. Given the nature of Shot’s microtubule-actin crosslinking activity, it could serve as an organizer of microtubules along the actin filaments of the ring canals on the nurse cell side and facilitate their transport towards the oocyte.

### The “go-with-the-flow” mechanism of cytoplasmic transport

Here we report that the minus-end directed motor, cytoplasmic dynein, glides microtubules, and microtubules in turn stir the cytoplasm and transfer cytoplasmic contents from nurse cells to the oocyte. To our knowledge, this is the first report of microtubule gliding by cortical dynein driving cargo movement in interphase cells (the “go-with-the-flow” mechanism).

Previously, we have demonstrated that conventional kinesin, kinesin-1, a major microtubule plus-end motor, can slide microtubules against each other ^27, 28, 61–63^, and this microtubule sliding drives ooplasmic streaming, bulk circulation of the entire cytoplasm in late-stage oocytes that is essential for localization of the posterior determinant, *osk*/Staufen RNPs ^29, 52^.Thus, both major microtubule motors, plus-end directed kinesin-1 and minus-end directed dynein, in addition to canonical cargo transport along microtubules, can drive bulk transport of viscous cytoplasm. Yet kinesin-1 and dynein drive bulk movement in different manners: kinesin-1 drives microtubule sliding against each other, while dynein glides microtubules along the cortex; and for different purposes: kinesin-1 powers intracellular circulation, whereas dynein propels intercellular transport.

This “go-with-the-flow” mechanism is highly efficient for cargo delivery, especially in large cells, such as the oocyte. The dynein-driven cytoplasmic flow allows the oocyte to acquire cytoplasmic materials for its rapid growth. Interestingly, the ring canals, have been observed in female germline cells of vertebrate organisms (e.g., human, rabbit, rat, hamster, mouse, chicken, and frog) ^64^. Particularly, it has been shown that cytoplasmic contents such as mitochondria and Golgi material are transferred to the mouse oocyte from interconnected cyst cells in a microtubule-dependent fashion ^65^ (Niu W. and Spradling AC. 2021. Mouse oocytes develop in cysts with the help of nurse cells. bioRxiv. doi: https://doi.org/10.1101/2021.11.04.467284). As dynein is highly conserved across species, it is important to have further studies to examine whether it plays a similar role in germline cytoplasmic transfer in higher organisms.

## Materials and Methods

### *Drosophila* strains

Fly stocks and crosses were maintained on standard cornmeal food (Nutri-Fly® Bloomington Formulation, Genesee, Cat #: 66-121) supplemented with dry active yeast at room temperature (∼24– 25°C). The following fly stocks were used in this study: *mat αtub-Gal4^[V37]^* (III, Bloomington *Drosophila* Stock Center #7063); *Act5C-Gal4* (III, Bloomington *Drosophila* Stock Center #3954); *nos-Gal4-VP16* (III, from Dr. Edwin Ferguson, the University of Chicago ^66, 67^); *UAS-Dhc64C-RNAi* (line #1: TRiP.GL00543, attP40, II, Bloomington *Drosophila* Stock Center #36583, targeting DHC64C CDS 10044–10064 nt, 5’-TCGAGAGAAGATGAAGTCCAA-3’; line #2: TRiP.HMS01587, attP2, III, Bloomington *Drosophila* Stock Center #36698, targeting DHC64C CDS 1302–1322 nt, 5’-CCGAGACATTGTGAAGAAGAA-3’) ^28, 68^; *UASp-Gl^ΔC^* (1-826 residues of DCTN1/p150^Glued^, based on the *DCTN1/p150*^1^ mutation) (II, 16.1, from Dr. Thomas Hays, University of Minnesota) ^12^; *UAS-Lis1-RNAi* (II, from Dr. Graydon Gonsalvez, Augusta University, targeting Lis1 CDS 1197-1217 nt, 5’-TAGCGTAGATCAAACAGTAAA-3’) ^19^; *UAS-BicD-RNAi* (TRiP.GL00325, attP2, III, Bloomington *Drosophila* Stock Center #35405, targeting BicD 3’UTR 639-659 nt, 5’-ACGATTCAGATAGATGATGAA-3’); *UAS-Spindly-RNAi* (TRiP.HMS01283, attP2, III, Bloomington *Drosophila* Stock Center #34933, targeting Spindly CDS 1615–1635 nt, 5’-CAGGACGCGGTTGATATCAAA-3’) ^56^; *UAS-hook-RNAi* (TRiP.HMC05698, attP40, II, Bloomington *Drosophila* Stock Center #64663, targeting HOOK CDS 241-261 nt, 5’-TACGACTACTACAGCGACGTA-3’); *UAS-GFP-RNAi* (Bloomington *Drosophila* Stock Center #41551); *UASp-tdMaple3-αtub84*B (II) ^29^; *UASp-EMTB-3XTagRFP* (III) ^30^; *UASp-GFP-Patronin* (II) (from Dr. Uri Abdu, Ben-Gurion University of the Negev) ^30, 52, 69^; *Jupiter-GFP* (protein trap line ZCL2183, III) ^27, 28, 31^; *UASp-LifeAct-TagRFP* (III, 68E, Bloomington *Drosophila* Stock Center # 58714); *UASp-F-Tractin-tdTomato* (II, Bloomington *Drosophila* stock center #58989) ^43^; *UAS-Zipper-RNAi* (TRiP.GL00623, attP40, II, Bloomington *Drosophila* Stock Center #37480, targeting Zipper 3’UTR 36-56 nt, 5’-CAGGAAGAAGGTGATGATGAA-3’); *hs-FLP^[^*^12^*^]^* (X, Bloomington *Drosophila* Stock Center #1929); *FRTG13 ubi-GFP.nls* (II, Bloomington *Drosophila* Stock Center # 5826); *FRTG13 zip*^2^/*CyO* (Bloomington *Drosophila* stock center # 8739); *sqh-GFP-RLC* (III, Bloomington *Drosophila* stock center #57145); *UASp-Mito-MoxMaple3* (II) ^15^; *ubi-GFP-Pav* (II, from Dr. David Glover, Caltech) ^70^; *UASp-RFP-Golgi* (II, Bloomington *Drosophila* Stock Center # 30908, aka *UASp-GalT-RFP*) ^71^; *pDlic-Dlic-GFP* (II, under the control of its native promoter, from Dr. Thomas Hays, University of Minnesota) (Neisch et al., submitted, reagent shared pre-publication). The following fly stocks were generated in this study using either PhiC31-mediated integration or P-element transformation: *UASp-Dlic-RNAi* (targeting Dlic 3’UTR 401-421 nt, 5’-AGAAATTTAACAAAAAAAAAA -3’, in pWalium22 vector, inserted at attP-9A (VK00005) 75A10 site, III, M5)*; UASp-GEM* (III, M1)*; UASp-F-Tractin-Myc-BicD* (II, M4)*; UASp-F-Tractin-GFP-Kin14VIb* (II, M1); *UASp-Myc-HA-DlicNT-LOV2 (M2) at attP14 (36A10); UASp-Zdk1-DlicCT-sfGFP-Myc (M2) at attP33 (50B6)*.

### Plasmid constructs

#### pWalium22-Dlic3’UTR-shRNA

The oligos of Dlic3’UTR-shRNA (agt<u>AGAAATTTAACAAAAAAAAAA</u>tagttatattcaagcata<u>TTTTTTTTTTGTTAAATTTCT</u>gc) were synthesized and inserted into the pWalium22 vector (*Drosophila* Genomics Resource Center, Stock Number #1473, 10XUAS) ^72^ by NheI(5’)/EcoRI(3’).

#### pUASp-GEM

pCDNA3.1-pCMV-PfV-GS-Sapphire was a gift from Liam Holt (Addgene plasmid # 116933; RRID:Addgene_116933) ^42^. GEM (PfV-GS-Sapphire) was cut and inserted into the pUASp vector by NheI(5’)/XbaI(3’).

#### pUASp-F-Tractin-Myc-BicD

F-Tractin (the actin binding domain of rat Inositol trisphosphate 3-kinase A (lTPKA), residues 9-40, atgGGCATGGCGCGACCACGGGGCGCGGGGCCCTGCAGCCCCGGGTTGGAGCGG GCTCCGCGCCGGAGCGTCGGGGAGCTGCGCCTGCTCTTCGAA) ^43, 73^ was synthesized and inserted into pUASp by KpnI (5’)/SpeI (3’). Myc-BicD was amplified from pAC-FRB-GFP-BicD^68^ and inserted into pUASp vector by SpeI (5’)/XbaI (3’).

#### pUASp-F-Tractin-GFP-Kin14VIb

F-Tractin (the actin binding domain of rat Inositol trisphosphate 3-kinase A (lTPKA), residues 9-40, atgGGCATGGCGCGACCACGGGGCGCGGGGCCCTGCAGCCCCGGGTTGGAGCGG GCTCCGCGCCGGAGCGTCGGGGAGCTGCGCCTGCTCTTCGAA) ^43, 73^ was synthesized and inserted into pUASp by KpnI (5’)/SpeI (3’). EGFP and Kin14VIb were amplified from the vector of pET15b-EGFP-GCN4-kinesin14VIb (a gift from Dr. Gohta Goshima) ^44^ by PCR and inserted into pUASp by SpeI(5’)/NotI(3’) and NotI(5’)/XbaI (3’), respectively.

#### pUASp-Myc-HA-DlicNT-LOV2-attB

DlicNT (1-370 residues) and LOV2[WT] were amplified by PCR from pAC-Dlic-EGFP (a gift from Dr. Thomas Hays, University of Minnesota) ^74^ and pTriEx-NTOM20-mVenus-LOV2[WT] (a gift from Dr. Klaus Hahn, University of North Carolina at Chapel Hill) ^48^ and inserted into pMT.A vector by EcoRV(5’)/XhoI(3’) and XhoI(5’)/XbaI(3’), respectively. Myc-HA with linkers (GGSG) was synthesized and inserted into the pMT-A-DlicNT-LOV2[WT] by EcoRI(5’)/EcoRV(3’). Myc-HA-DlicNT-LOV2[WT] was subcloned from the pMT.A-Myc-HA-DlicNT-LOV2[WT] into the pUASp vector by KpnI(5’)/XbaI(3’), and attB site was subcloned from the pUASp-attB vector (*Drosophila* Genomics Resource Center/DGRC vector #1358) by AatII(5’)/AatII(3’) to create pUASp-Myc-HA-DlicNT-LOV2-attB.

#### pUASp-Zdk1-DlicCT-sfGFP-Myc-attB

Zdk1 and C-terminus of Dlic (371-493 residues) were amplified by PCR from pTriEx-mCherry-Zdk1 (a gift from Dr. Klaus Hahn, University of North Carolina at Chapel Hill) ^48^) and pAC-Dlic-EGFP (a gift from Dr. Thomas Hays, University of Minnesota) ^74^, and inserted into pMT.A by SpeI(5’)/EcoRI(3’) and EcoRI(5’)/EcoRV(3’), respectively. Superfolder GFP (sfGFP) with a Myc tag was inserted into the pMT.A-Zdk1-DlicCT by EcoRV(5’)/XbaI(3’). Zdk1-DlicCT-sfGFP-myc was then subcloned into the pUASp-attB vector (*Drosophila* Genomics Resource Center/DGRC vector #1358) by SpeI(5’)/XbaI(3’) to create pUASp-Zdk1-DlicCT-sfGFP-Myc-attB.

#### pcDNA3.1(+)-Myc-HA-DlicNT-LOV2

Myc-HA-DlicNT-LOV2 was subcloned from pMT.A-Myc-HA-DlicNT-LOV2[WT] into pcDNA3.1(+) by KpnI(5’)/XbaI(3’).

#### pcDNA3.1(+)-Zdk1-DLicCT-sfGFP-Myc

Zdk1-DlicCT-sfGFP-Myc was amplified by PCR from pMT.A-Zdk1-DlicCT-sfGFP-Myc and inserted into pcDNA3.1(+) by HindIII(5’)/XbaI(3’).

### Immunostaining of *Drosophila* egg chambers

A standard fixation and staining protocol was previously described ^15, 29, 30, 52, 67^. Samples were stained with primary antibody at 4°C overnight and with fluorophore-conjugated secondary antibody at room temperature (24∼25°C) for 4 h. Primary antibody used in this study: mouse monoclonal anti-Orb antibody (Orb 4H8, Developmental Studies Hybridoma Bank, supernatant, 1:5); rabbit phospho-MLC (pMLC 2/Ser19, 1:100, Cell Signaling, Cat# 3671); mouse monoclonal anti-DHC antibody (2C11-2, Developmental Studies Hybridoma Bank, concentrate, 1:50); mouse monoclonal anti-Myc antibody (1-9E10.2, 1:100) ^75^.

Secondary antibody used in this study: FITC-conjugated or TRITC-conjugated anti-mouse secondary antibody (Jackson ImmunoResearch Laboratories, Inc; Cat# 115-095-062 and Cat# 115-025-003) at 10 µg/ml; FITC-conjugated anti-rabbit secondary antibody (Jackson ImmunoResearch Laboratories, Inc; Cat#111-095-003) at 10 µg/ml.

Some samples were stained with rhodamine-conjugated phalloidin (0.2 µg/ml) and DAPI (1 µg/mL) for 1 ∼2 h before mounting. Samples were imaged on a Nikon A1plus scanning confocal microscope with a GaAsP detector and a 20× 0.75 N.A. lens using Galvano scanning, a Nikon W1 spinning disk confocal microscope (Yokogawa CSU with pinhole size 50 µm) with Photometrics Prime 95B sCMOS Camera or Hamamatsu ORCA-Fusion Digital CMOS Camera and a 40 x 1.30 N.A. oil lens or a 40X 1.25 N.A. silicone oil lens, or a Nikon Eclipse U2000 inverted stand with a Yokogawa CSU10 spinning disk confocal head with an Photometrics Evolve EMCCD camera and a 40× 1.30 N.A. oil lens, all controlled by Nikon Elements software. Z-stack.images were acquired every 1 µm/step for whole ovariole imaging or 0.3∼0.5 µm/step for individual egg chambers.

### Microtubule staining in *Drosophila* egg chambers

Ovaries were dissected in 1XPBS and fixed in 8% EM-grade formaldehyde +1X Brinkley Renaturing Buffer 80 (BRB80, 80 mM piperazine-N,N’-bis(2-ethanesulfonic acid) [PIPES]), 1 mM MgCl2, 1 mM EGTA, pH 6.8) +0.1%Triton X-100 for 20 min on the rotator; briefly washed with 1XPBTB (1XPBS+0.1% Triton X-100+0.2% BSA) five times and stained with FITC-conjugated β-tubulin antibody (ProteinTech, Cat# CL488-66240) 1:100 at 4C overnight; then samples were stained rhodamine-conjugated phalloidin and DAPI for 1 h before mounting. Samples were imaged using Nikon W1 spinning disk confocal microscope (Yokogawa CSU with pinhole size 50 µm) with Photometrics Prime 95B sCMOS Camera, and a 40X 1.25 N.A. silicone oil lens, controlled by Nikon Elements software. Images were acquired every 0.3 µm/step in z stacks and 3D deconvolved using Richardson-Lucy iterative algorithm provided by Nikon Elements.

### Live imaging of *Drosophila* egg chamber

Young mated female adults were fed with dry active yeast for 16∼18 hours and then dissected in Halocarbon oil 700 (Sigma-Aldrich, Cat# H8898) as previously described ^15, 29, 30, 52^. Fluorescent samples were imaged using Nikon W1 spinning disk confocal microscope (Yokogawa CSU with pinhole size 50 µm) with Photometrics Prime 95B sCMOS Camera or Hamamatsu ORCA-Fusion Digital CMOS Camera, and a 40 x 1.30 N.A. oil lens or a 40X 1.25 N.A. silicone oil lens, controlled by Nikon Elements software.

### Photoconversion of tdMaple3-tubulin and Mito-MoxMaple3 in ovary

Photoconversions of tdMaple3-tubulin and Mito-MoxMaple3 were performed using illumination from a Heliophor 89 North light in the epifluorescence pathway by a 405 nm filter, either locally (for tdMaple3-tubulin) or globally (for Mito-MoxMaple3) through an adjustable pinhole in the field diaphragm position for 10∼20 s. Samples were imaged either on a Nikon Eclipse U2000 inverted stand with a Yokogawa CSU10 spinning disk confocal head with an Photometrics Evolve EMCCD camera and a 40× 1.30 N.A. oil lens (for tdMaple3-tubulin), or a Nikon W1 spinning disk confocal microscope (Yokogawa CSU with pinhole size 50 µm) with Hamamatsu ORCA-Fusion Digital CMOS Camera and a 40X 1.25 N.A. silicone oil lens (Mito-MoxMaple3), all controlled by Nikon Elements software.

### Induction of *zip*^2^ germline clones

*FRTG13 zip*^2^*/CyO* virgin female flies were crossed with males carrying *hs-flp^[^*^12^*^]^/y; FRTG13 ubi-GFP.nls/CyO*. From the cross, young pupae at day 7 and day 8 AEL (after egg laying) were subjected to heat shock at 37 °C for 2 hours each day. Non CyO F1 females were collected 3-4 day after heat shock and fattened with dry active yeast overnight before dissection for fixation and Orb staining.

### Extraction and fixation of ovary samples

Ovaries were dissected and gently teased apart in 1X Brinkley Renaturing Buffer 80 (BRB80, 80 mM piperazine-N,N’-bis(2-ethanesulfonic acid) [PIPES]), 1 mM MgCl2, 1 mM EGTA, pH 6.8). The dissected samples were extracted in 1X BRB80+1% Triton X-100 for 20 min without agitation. After the extraction, the samples were fixed with EM-grade formaldehyde in 1X BRB80+0.1% Triton X-100 for 20min on rotator, washed with 1XPBTB (1XPBS+0.1% Triton X-100+0.2% BSA) five times before immunostaining.

### Measurement of nurse cell size

Z stacks of triple color images of the ovarioles from *yw; ubi-GFP-Pav* stained with rhodamine conjugated-phalloidin and DAPI were acquired, and nurse cell area was specified (at the largest cross-section) and measured by manual polygon selection (area size) in FIJI ^76^.

### S2R+ cell transfection and imaging acquisition

*Drosophila* S2R+ cells were transfected with 0.5 µg DNA (pAC-Gal4+pUASp-GEM, or pAC-LAMP1-GFP ^77^) in 12-well plate using Effecetene transfection kit (Qiagen, Cat. # / ID: 301425) and then were plated 48 hours after transfection on concanavalin A-coated glass coverslips ^78^. Cells were imaged using Nikon W1 spinning disk confocal microscope (Yokogawa CSU with pinhole size 50 µm) with Photometrics Prime 95B sCMOS Camera, and a 100 x 1.45 N.A. oil lens, controlled by Nikon Elements software.

### Measurement of GEM particle movement in control and dynein-RNAi

Samples expressing GEM particles (*yw; mat atub-Gal4^[V37]^*/*UASp-GEM* and *yw; UAS-Dhc64C-RNAi/+; mat atub-Gal4^[V37]^/UASp-GEM*) were imaged on a Nikon W1 spinning disk confocal microscope (Yokogawa CSU with pinhole size 50 µm) with Photometrics Prime 95B sCMOS Camera and a 40X 1.25 N.A. silicone oil lens for 2 min at the frame rate of every 2 sec, controlled by Nikon Elements software. Images were processed in Fiji and analyzed DiaTrack 3.04 Pro ^79^, with a maximum particle jump distance of 3.2 μm/s.

### HEK293 cell transfection and GFP-binder pull-down assay

HEK293 cells were co-transfected with pcDNA3.1(+)-myc-HA-DlicNT-LOV2 and pcDNA3.1(+)-Zdk1-DlicCT-sfGFP-Myc by Calcium Phosphate transfection as previously described ^80^. 40∼48 hrs later cell extracts from transfected cells were made with red-only light on (no blue light, considered as “dark”): cells were rinsed twice with 1XPBS and scraped into 350 ul extraction buffer (10mM Tris-buffer, ph7.4, 50mM NaCl; 0.5 mM MgCl2); cracked in cell cracker and collected in 1.5 ml microtube. Triton X-100 was added to the extract to final concentration of 1%. The extract was centrifuged at 14k for 15 min at 4°C (Eppendorf, 5804R). Supernatant was collected and divided equally in two parts. Both samples were mixed with 25 µl pre-washed GFP-binder agarose beads (Chromotek) ^81^. One sample was kept in aluminum foil for 2 hours at 4C (dark); another sample was illuminated first with blue light for 1 min and then full natural light for 2 hours at 4°C. Beads were washed twice, and eluted with 40-50 µl 5x sample buffer. Initial cell extract and both eluted samples were separated by electrophoresis in 8% SDS-PAGE gel, and transferred to nitrocellulose membrane (Odyssey Nitrocellulose Membrane, LI-COR). Membrane was blotted with mouse monoclonal anti-Myc antibody (1-9E10.2, 1:1000), followed by HRP-conjugated goat-anti-mouse secondary antibody (Jackson ImmunoResearch Laboratories, Cat# 115-035-003, 1:10,000). The blot was developed on Odyssey phosphoimager (LI-COR).

### Statistical analysis

The plots in figures show either percentage of phenotypes, or average values, as indicated in figure legends. Error bars represent 95% confidence intervals. N stands for number of samples examined in each assay, unless it is specified elsewhere in figure legends. Unpaired t tests with Welch’s correction were performed in GraphPad Prism 8.0.2. P values and levels of significance are listed in figure legends.

## Acknowledgements

We would like to thank many colleagues who generously contributed reagents for this work: Dr. Thomas Hays (University of Minnesota); Dr. Graydon Gonsalvez (Augusta University); Dr. Uri Abdu (Ben-Gurion University of the Negev, Israel); Dr. David Glover (Caltech); Dr. Gohta Goshima (Nagoya University, Japan); Dr. Klaus Hahn (University of North Carolina Chapel Hill); Dr. Andrew Carter (MRC Laboratory of Molecular Biology, Cambridge); Dr. Edwin Ferguson (the University of Chicago); Bloomington *Drosophila* Stock Center (supported by National Institutes of Health grant P40OD018537) and *Drosophila* Genomics Resource Center (supported by NIH grant 2P40OD010949). The Orb 4H8 monoclonal antibody developed by Dr. Paul D. Schedl’s group at Princeton University and DHC 2C11-2 antibody developed by Dr. Jonathan M. Scholey’s group at University of California, Davis, were obtained from the Developmental Studies Hybridoma Bank, created by the NICHD of the NIH and maintained at The University of Iowa, Department of Biology, Iowa City, IA 52242. We also thank all the Gelfand laboratory members for support, discussions, and suggestions.

Research reported in this study was supported by the National Institute of General Medical Sciences grants R01GM124029 and R35GM131752 to V.I. Gelfand.

## Author contributions

W.L. and V.I.G. planned and designed the research. W.L., M.L., A.S.S. and V.I.G. conducted experiments and data analysis; W.L. and V.I.G. wrote the manuscript.

## Data Availability and Code Availability statement

All data generated and analyzed during this study are included in this manuscript. This study didn’t generate computational code.

## Supplementary figures and figure legends

**Supplementary Figure 1.**
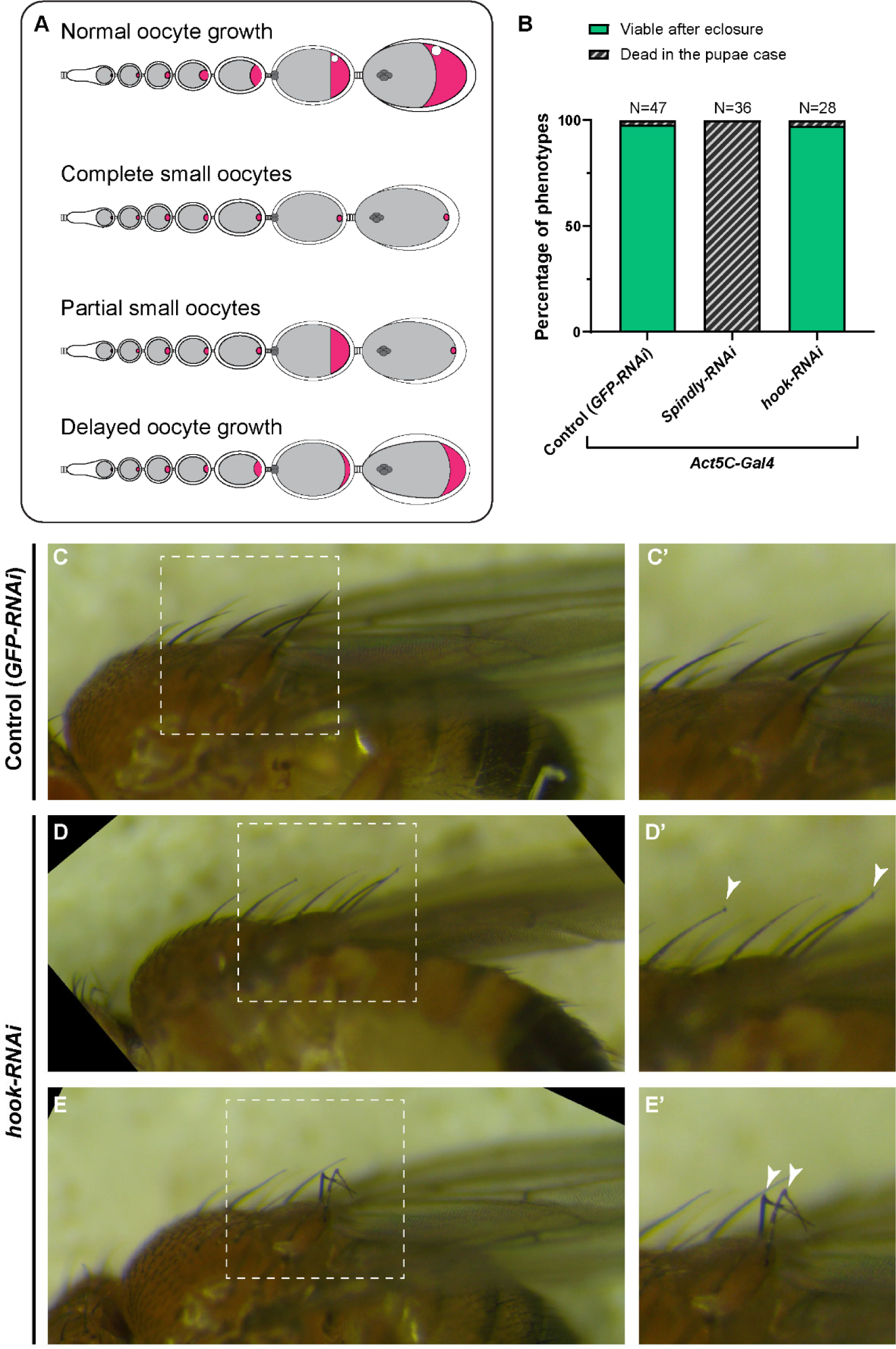
Summary of dynein-related RNAi phenotypes. Related to Figure 1. (A) A Cartoon illustration of oocyte growth phenotypes (oocytes are highlighted in magenta): (1) normal oocyte growth: all egg chambers in the single ovariole have normal-sized oocytes according to the stages; (2) complete small oocytes: oocyte growth is completely arrested throughout the stages, (3) partial small oocyte growth: some egg chambers display oocyte growth arrest, while others have normal-size oocytes in the same ovariole; (4) delayed oocyte growth: oocyte size is smaller compared to wild-type at certain stages. (B) Viability assay (from pupae to adults) in listed genotypes. As the *Act5C-Gal4* transgene is balanced with a TM6B (Tb) balancer, only non-Tb pupae were selected for this assay. (C-E’) Bristle phenotypes in control (C) and in *hook-RNAi* (D-E) male adults. Control and *hook-RNAi* are of the same genotypes as (B). Mild (D) and severe (E) hooked bristle phenotypes are seen in *Act5C>hook-RNAi* flies.

**Supplementary Figure 2.**
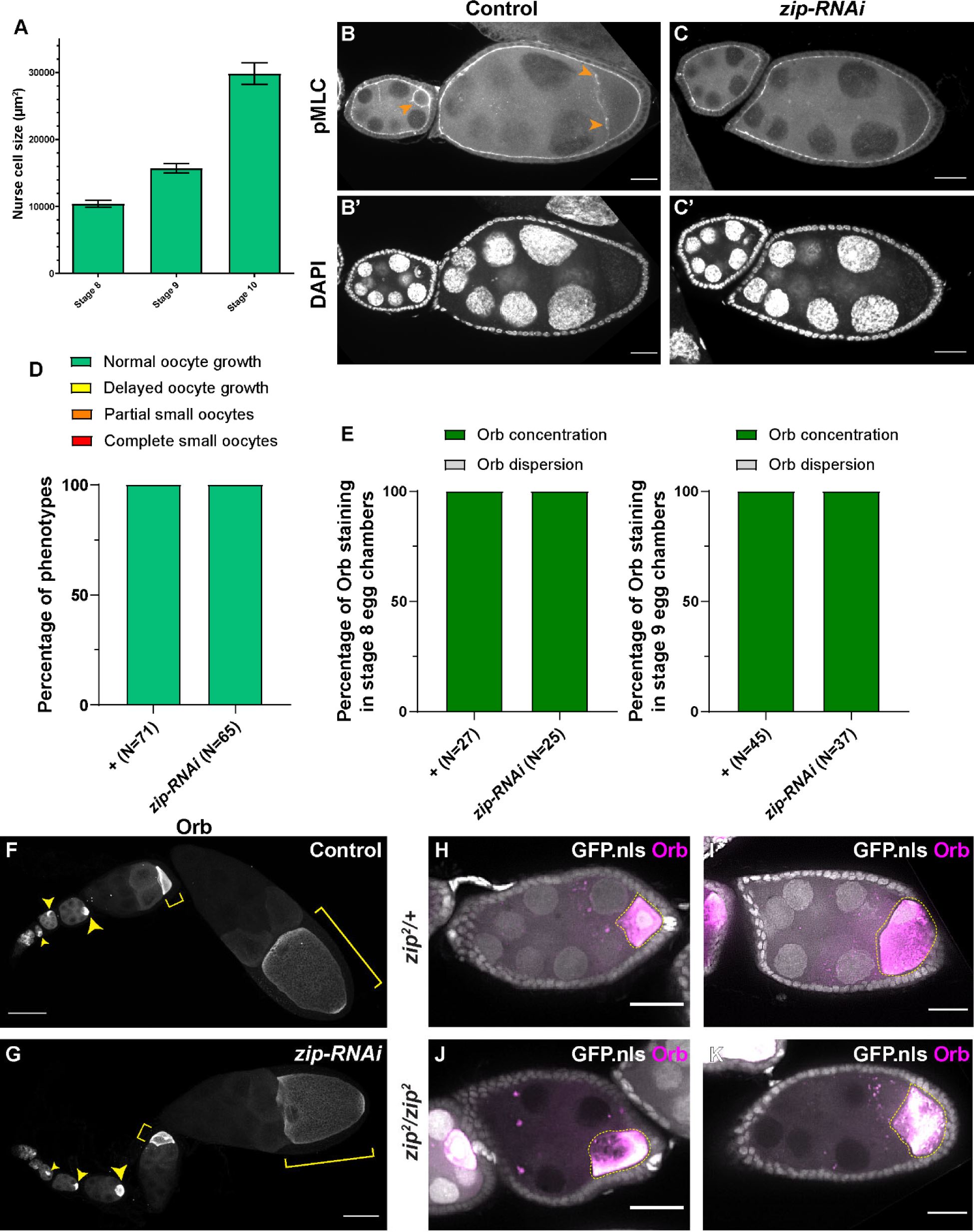
Myosin-II activity is dispensable for oocyte growth during mid-oogenesis. Related to Figure 2. (A) Nurse cell size in stage 8 to stage 10 egg chambers. Stage 8, N=13; stage 9, N=23; stage 10, N=14. (B-C’) Phospho-myosin-II light chain (pMLC) antibody staining in control and in *zip-RNAi* samples. The pMLC staining between the nurse cells and the oocytes is abolished in *zip-RNAi* (C), compared to control (B, pointed with orange arrowheads). To note, the pMLC staining at the apical side of the somatic follicle cells are not affected by the germline knockdown of *zip-RNAi* and servers as an internal control of the antibody staining. Scale bars, 20 µm. (D-E) Summaries of oocyte growth phenotypes (D) and Orb staining phenotypes (E) in control and in *zip-RNAi*. Both control and *zip-RNAi* carry one copy of *maternal αtub-_Gal4_[V37]*. (F-G) Characteristic Orb staining in control (F) and *zip-RNAi* (G) ovarioles. Oocytes are highlighted with either yellow arrowheads or yellow brackets. Scale bars, 50µm. Both samples carry one copy of *maternal αtub-Gal4^[V^*^37^*^]^*. (H-K) Orb staining in *zip*^2^ heterozygous (H-I) and homozygous (J-K, marked with the absence of nuclear-localized GFP signal) egg chambers. Among 46 *zip*^2^ homozygous egg chambers examined, 28 egg chambers have proper oocyte specification (shown by Orb concentration). 25 out 28 *zip*^2^ homozygous egg chambers display no major growth defects in mid-oogenesis. To note: Orb staining becomes chunky and less evenly localized in the *zip*^2^ homozygous oocytes. Scale bars, 50 µm.

**Supplementary Figure 3.**
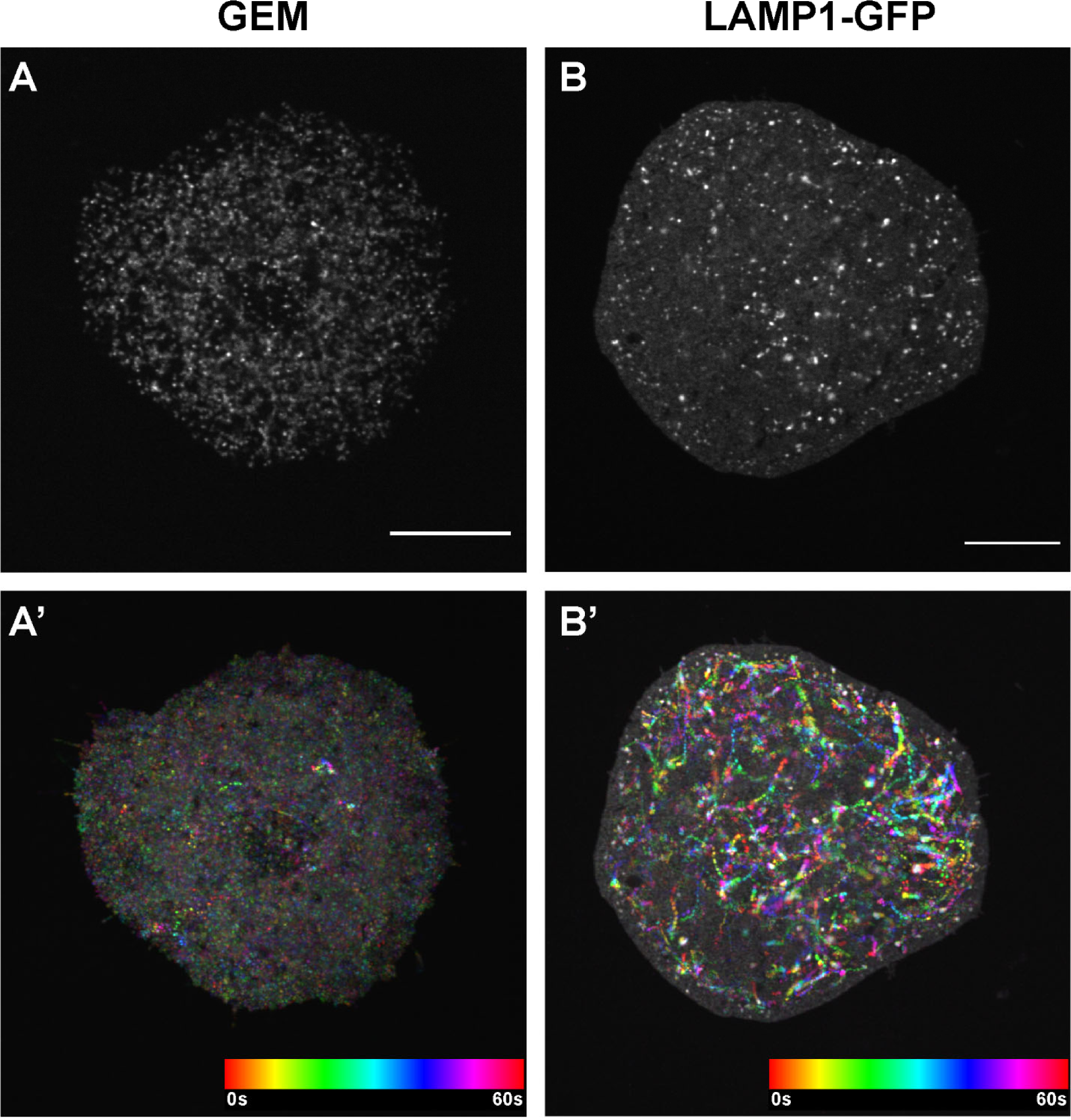
GEMs and lysosomes in *Drosophila* S2R+ cells. Related to Figure 3. Single frame images (A-B) and temporal color-coded hyperstacks (A’-B’) of GEM particles (labeled with pAC-Gal4+pUASp-GEM) (A-A’) and lysosomes (labeled with pAC-LAMP1-GFP) (B-B’) in *Drosophila* S2R+ cells on Concanavalin-A coated coverslips. Scale bars, 10 µm.

**Supplementary Figure 4.**
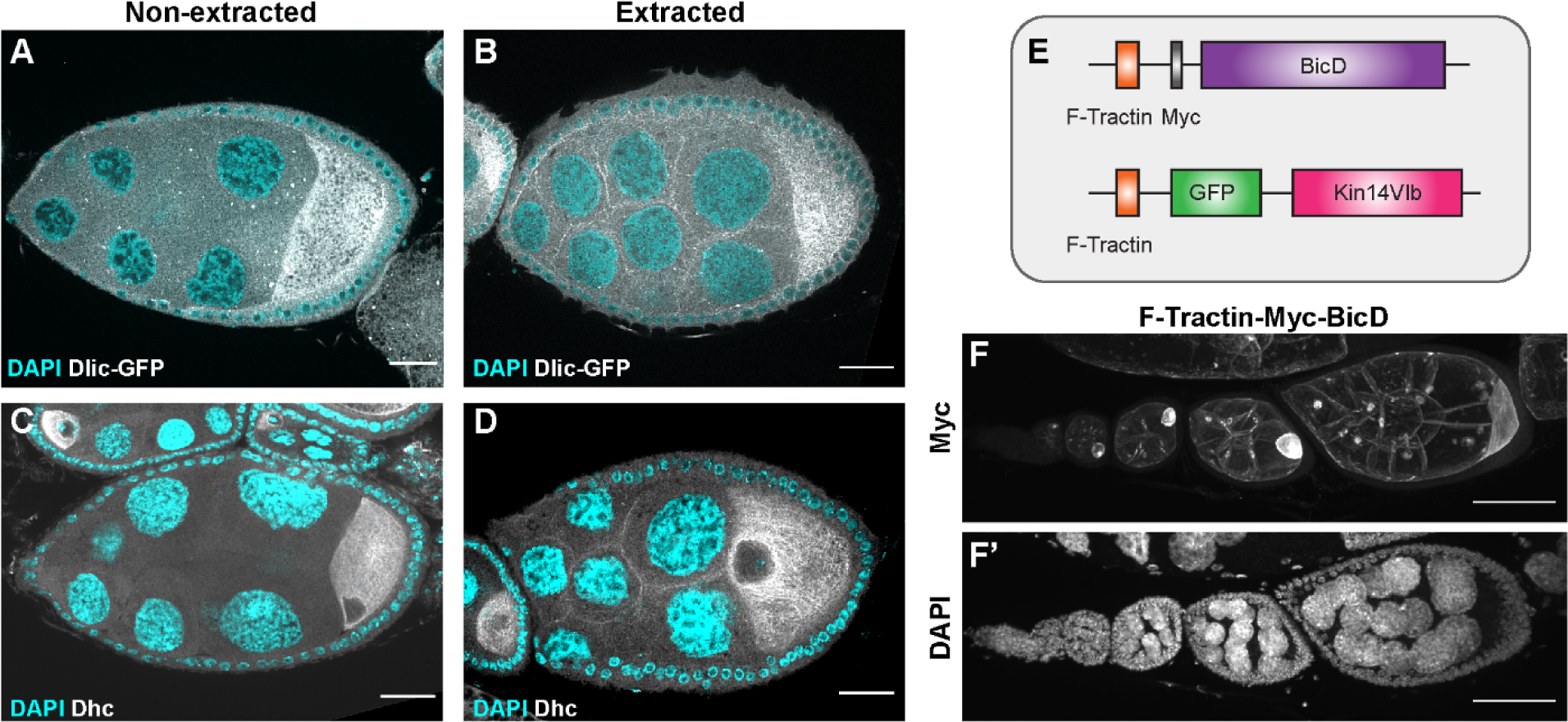
The recruitment of dynein to nurse cell cortex. Related to Figure 4. (A-D) Dlic-GFP under its endogenous promoter (A-B) and DHC antibody staining (C-D) in non-extracted (A, C) and extracted (B, D) samples. After extraction, both Dlic-GFP and Dhc staining show clear cortical localization in nurse cells. Scale bars, 20 µm. (E) A schematic illustration of the cortically-recruited BicD and kin14 constructs. (F-F’) F-Tractin-Myc-BicD is predominantly localized to the cell cortex. The expression is driven by one copy of *maternal αtub-Gal4^[V^*^37^*^]^*. Scale bars, 50 µm.

**Supplementary Figure 5.**
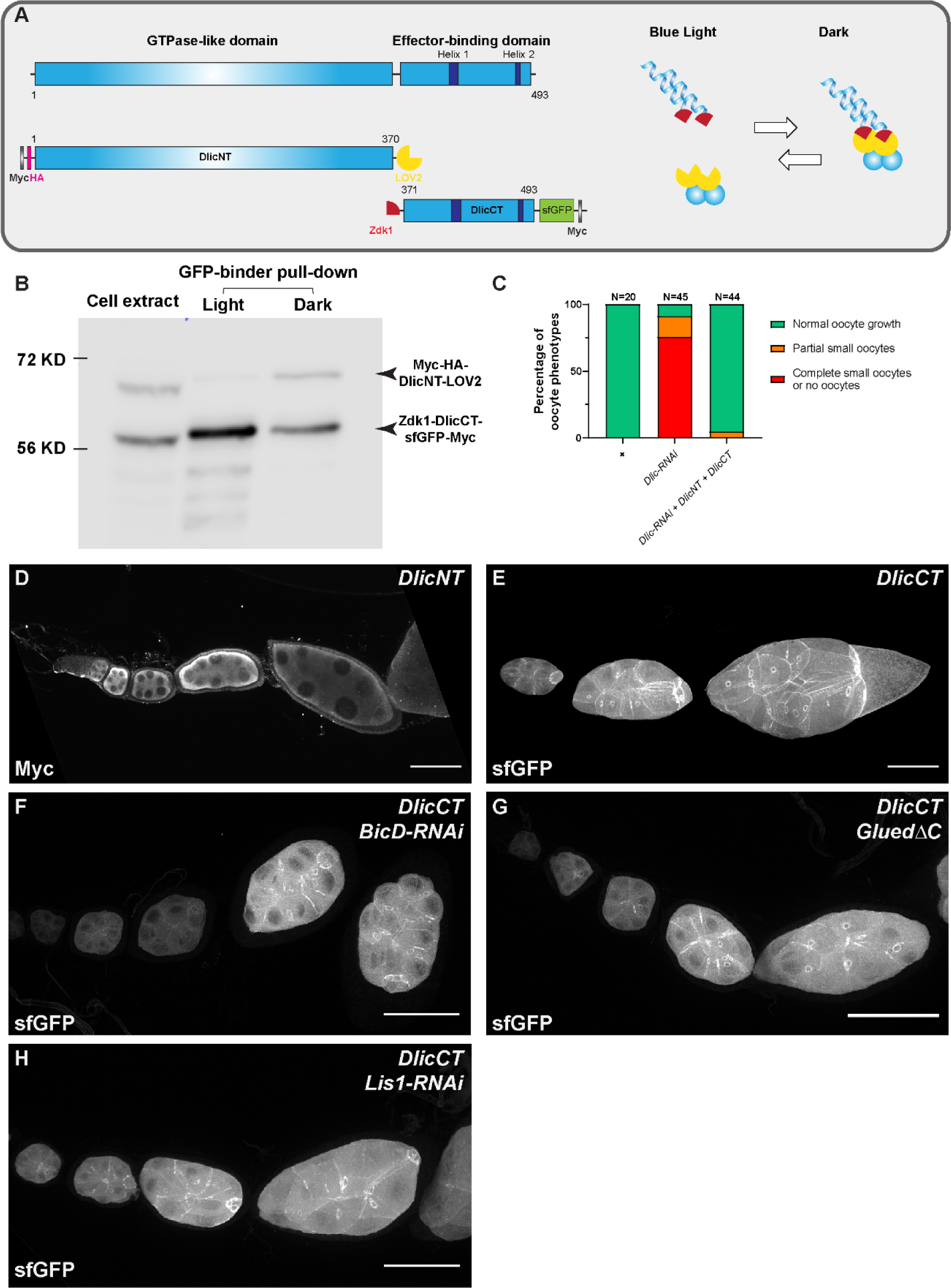
Dlic C-terminus is sufficient for targeting to nurse cell cortex. Related to Figure 5. (A) A schematic illustration of *Drosophila* Dlic and Dlic truncations (DlicNT and DlicCT). C-terminus of DlicNT and N-terminus of DlicCT are tagged with the LOVTRAP probes, LOV2 and Zdk1, respectively. LOV2 interacts with Zdk11 in dark, and dissociates from Zdk1 in the presence of blue light. (B) Zdk1-DlicCT interacts with DlicNT-LOV2 in dark. GFP-binder was used to pull down Zdk1-DlicCT in cell extracts from HEK293 cells expressing both DlicNT and DlicCT constructs, and anti-Myc antibody was used to probe for Dlic truncations. Lane 1: cell extract; Lane 2: GFP-binder pulldown sample in light; lane 3; GFP-binder pulldown sample in dark. (C) Coexpression of DlicNT and DlicCT reuses the oogenesis defects caused by *Dlic-RNAi*. The expression is driven by one copy of *nos-Gal4^[VP^*^16^*^]^*. (D) DlicNT localization in the germ line. The expression is driven by one copy of *maternal αtub-Gal4^[V^*^37^*^]^*. A small Z-projection (∼5 µm) is used to show the DlicNT localization. To note: overexpression of DlicNT driven by *maternal αtub-Gal4^[V^*^37^*^]^* results in delayed oocyte growth. Scale bar, 50 µm. (E-H) DlicCT localizations in control (E), *BicD-RNAi* (F), Dynactin/p150 inhibited (*GluedΔC*, G) and *Lis1-RNAi* (H) egg chambers. Whole ovariole z-projections (30 ∼ 40 µm) are used to show the DlicCT cortical localization. The numbers of ovarioles examined: Control, N=28; *BicD-RNAi*, N=39; *GluedΔC*, N=30; *Lis1-RNAi*, N=29. All listed genotypes are with one copy of *maternal αtub-Gal4^[V^*^37^*^]^*. Scale bars, 50 µm.

## Videos

**Video 1.** Microtubule movement (labeled with locally photoconverted tdMaple3-αtub, red) in control nurse cells. Scale bars, 20 µm. Related to Figure 2.

**Video 2.** Microtubules labeled with ectopically-expressed TagRFP-tagged human MAP7/Ensconsin microtubule binding domain (EMTB-3XTagRFP) move in nurse cells and through the nurse cell-oocyte ring canal (labeled with GFP-Pav in magenta). Scale bar, 50 µm. Related to Figure 2.

**Video 3.** Microtubules labeled with ectopically-expressed GFP-tagged minus-end binding protein, Patronin (GFP-Patronin) move in nurse cells and from the nurse cell to the oocyte through the ring canal (labeled with LifeAct-TagRFP in magenta). Scale bar, 50 µm. Related to Figure 2.

**Video 4.** Microtubules labeled with a GFP-tagged endogenous microtubule-associated protein Jupiter (Jupiter-GFP) in nurse cells and in the nurse cell-oocyte ring canal (labeled with F-Tractin-tdTomato in magenta). Scale bar, 50 µm. Related to Figure 2.

**Video 5.** No microtubule movement (labeled with locally photoconverted tdMaple3-αtub, red) in *Dhc64C-RNAi* nurse cells, compared to control. Scale bars, 20 µm. Related to Figure 2.

**Video 6.** Microtubule movement (labeled with a GFP-tagged endogenous MAP, Jupiter-GFP) in nurse cell-oocyte ring canals (labeled with GFP-Pav) in control, *zip-RNAi* and *Dhc64C-RNAi*. Scale bars, 10 µm. Related to Figure 2.

**Video 7.** Cytoplasmic flow from the nurse cells to the oocyte through ring canals (labeled with Sqh-GFP in magenta). Scale bar, 50 µm. Related to Figure 2.

**Video 8.** Bulk movement of mitochondria (labeled with Mito-MoxMaple3, without photoconversion, magenta) and Golgi units (labeled with RFP-Golgi, cyan) through the nurse cell-to-oocyte ring canal (labeled with GFP-Pav, magenta) in a stage 9 egg chamber. Scale bars, 50 µm. Related to Figure 2.

**Video 9.** Mitochondria movement (labeled with Mito-MoxMaple3, after global photoconversion, gray) in control, *zip-RNAi* and *Dhc64C-RNAi*. Ring canals are labeled with GFP-Pav (magenta); Scale bars, 50 µm. Related to Figure 2.

**Video 10.** Mitochondria movement (labeled with Mito-MoxMaple3, without photoconversion, left) and microtubule movement (labeled with EMTB-3XTagRFP, right) in nurse cell-oocyte ring canals (labeled with GFP-Pav, left). Scale bars, 50 µm in whole egg chambers, and 10 µm in zoom-in ring canals, respectively. Related to Figure 3.

**Video 11.** The movement of Golgi units (labeled with RFP-Golgi, left) and microtubules (labeled with Jupiter-GFP, right) in nurse cell-oocyte ring canals (labeled with F-Traction-tdTomato, left). Scale bars, 50 µm in whole egg chambers, and 10 µm in zoom-in ring canals, respectively. Related to Figure 3.

**Video 12.** GEM particles in a control egg chamber and in a *Dhc64C-RNAi* egg chamber. Scale bars, 20 µm. Related to Figure 3.

**Video 13.** GEM particles transport through a nurse cell-oocyte ring canal (labeled with F-Tractin-tdTomato, magenta) in control. Scale bar, 10 µm. Related to Figure 3.

**Video 14.** Mitochondria movement (labeled with Mito-MoxMaple3, after global photoconversion) in the nurse cell-oocyte ring canals of control, *Dlic-RNAi* and *DlicRNAi* + *F-*Tractin-GFP-Kin14VIb rescued samples. Ring canals are either labeled with GFP-Pav (in control and *Dlic-RNAi* samples) or F-Tractin-GFP-Kin14VIb (in the *Dlic-RNAi* + *F-*Tractin-GFP-Kin14VIb rescued sample), in magenta. Scale bars, 20 µm.

Related to Figure 4.

**Video 15.** Cortical dynein glides microtubules and transfers cytoplasm to the *Drosophila* oocyte. Related to Figure 5.

